# Decoupling simultaneous motor imagination and execution via orthogonal ECoG neural representations

**DOI:** 10.1101/2025.06.13.659445

**Authors:** Leonardo Pollina, Lucas Struber, Valeria de Seta, Eleonora Russo, Serpil Karakas, Stephan Chabardès, Tetiana Aksenova, Guillaume Charvet, Solaiman Shokur, Silvestro Micera

## Abstract

The brain coordinates multiple parallel motor programs, ensuring synergy and preventing interference during movements. Yet, performance often degrades when brain–machine interfaces are used during concurrent tasks or ongoing movements. We suggest that latent neural representations may represent a strategy to solve this issue. In this study, we addressed this question using neural signals from a tetraplegic individual with partial residual motor function, implanted with a wireless epidural electrocorticography (ECoG) device. By adapting dimensionality reduction techniques, we found that motor execution and motor imagery span partially overlapping subspaces in mesoscale neural signals, shaped by specific frequency band contributions. Despite substantial shared variance, we show that identifying orthogonal, condition-specific dimensions enables successful decoding of executed and imagined movements, even when performed simultaneously. These findings show that ECoG signals can expose separable neural subspaces, allowing executed and imagined actions to be harnessed independently and in concert. This opens a promising avenue to develop brain–machine interfaces that can simultaneously control multiple external devices or operate alongside natural movements.

## Introduction

Motor imagery is the cognitive process in which individuals mentally perform a movement without any physical execution. The relationship between motor execution and motor imagery has been extensively studied. Both engage motor-related brain regions but with different activation patterns [1–3]. Motor imagery is often considered a conscious parallel to motor preparation, where movements are mentally rehearsed before execution [4]. However, motor imagery also involves an inhibitory component consisting of the challenge of vividly imagining a movement while consciously suppressing its physical execution [5].

In the last decade, research has shifted from studying individual neurons to examining neural populations through large-scale recordings [6]. This shift has led to the concept of neural manifolds or neural spaces, lower-dimensional representations of neural activity that capture the coordinated dynamics of neural populations [7, 8]. These neural spaces provide valuable insights into how the brain encodes motor behaviors, even when motor processes activate corresponding brain areas without overt movements [9, 10]. The identification of neural spaces has shown that motor preparation and execution occupy distinct, orthogonal subspaces, with activity transitioning between these spaces as movements unfold [9, 11–13]. On the other hand, movement imagination [14] involves shared and exclusive spaces with motor execution, with the shared spaces consisting of common dynamics between the two neural processes and exclusive spaces containing neural patterns specific to each. While most of these studies rely on spike sorted or multi-unit activity, whether this geometric relationship between motor execution and imagery is also identifiable at the mesoscale level remains unclear. Addressing this question is particularly relevant for surface-electrode-based BMIs, such as electrocorticography (ECoG), which represent a compelling alternative to intracortical microelectrodes [15–18]. These BMIs offer reduced invasiveness and fewer associated side effects while preserving long-term usability [19–22]]. Identifying and leveraging neural spaces with these less invasive techniques could significantly enhance the effectiveness and accessibility of BMIs, broadening their impact for a wider range of patients. Moreover, the use of latent neural spaces both offers greater temporal stability, preserving a subject’s representation even as the activity of individual neurons varies over time [23–25], and allows the bridge across multiple subjects [26, 27], thereby enabling the training of more powerful and generalizable cross-subject BMI algorithms.

Most BMIs are designed to decode imagined movements in conditions with minimal or no concurrent physical movement. In certain instances, minor unstructured movements become part of the training signal distribution and are treated as noise by the classifier, thereby still enabling the decoding of motor imagery. While this strategy is appropriate for individuals with complete paralysis, it can be limiting for those with incomplete tetraplegia [28]. These individuals often retain some motor function, and restricting real movement in favor of motor imagery can hinder natural interaction in daily life. This highlights the need for alternative approaches that allow decoding of imagined actions concurrently with purposeful, goal-directed movements, whether coordinated with or independent from the external end effector controlled by the BMI. We propose that neural spaces capturing distinct representations of motor execution and motor imagery could help address this issue. Specifically, the separation of imagery and execution into orthogonal neural subspaces may help identify a specific neural space where motor imagery decoding could be more robust. This could ultimately allow users to perform physical movements without disrupting simultaneous BMI control through motor imagery. This challenge involves identifying task-null spaces, where neural signals can be modulated voluntarily without interfering with concurrent task-oriented actions [29]. By disentangling motor imagery from concurrent execution, we identify a candidate execution-null space, where neural modulation could drive an external effector, while performing other activities.

In this study, we addressed the challenge of decoupling motor execution from motor imagination by identifying a potential solution within the neural manifolds framework, which offers a principled approach to uncover condition-specific neural subspaces. To this end, we adapted dimensionality reduction methods originally developed for spiking activity to mesoscale cortical signals recorded from an individual with tetraplegia who retained some residual motor functions. Through this analysis, we identified partially overlapping neural representations of motor execution and imagination. Focusing on the neural subspaces exclusive to each condition, we found that they reliably captured condition-specific patterns with stable and robust dynamics. Using these representations, we were able to disentangle the neural processes underlying execution and imagination, even when they occurred simultaneously. Our findings indicate that this approach could enhance BMI performance by supporting user control of external interfaces via motor imagery alongside ongoing physical movements, without mutual interference.

## Results

A recent study has demonstrated that multi-unit neural activity related to motor execution and motor imagination overlap only partially in the neural space [14]. As depicted in **Fig.1A**, this suggests that the two activities share a common dimension (Dim. 1), while also maintaining exclusive dimensions, Dim. 2 for Execution and Dim. 3 for Imagination, respectively. We aimed to explore whether this relationship holds in neural spaces identified from mesoscale neural signals, such as those measured via ECoG. In this framework, we define a neural space, or neural manifold, as a space where the axes represent covariance patterns across channels and frequency bands, meaning that each axis represents a linear combination of these original features (**Fig.1B**).

**Figure 1:**
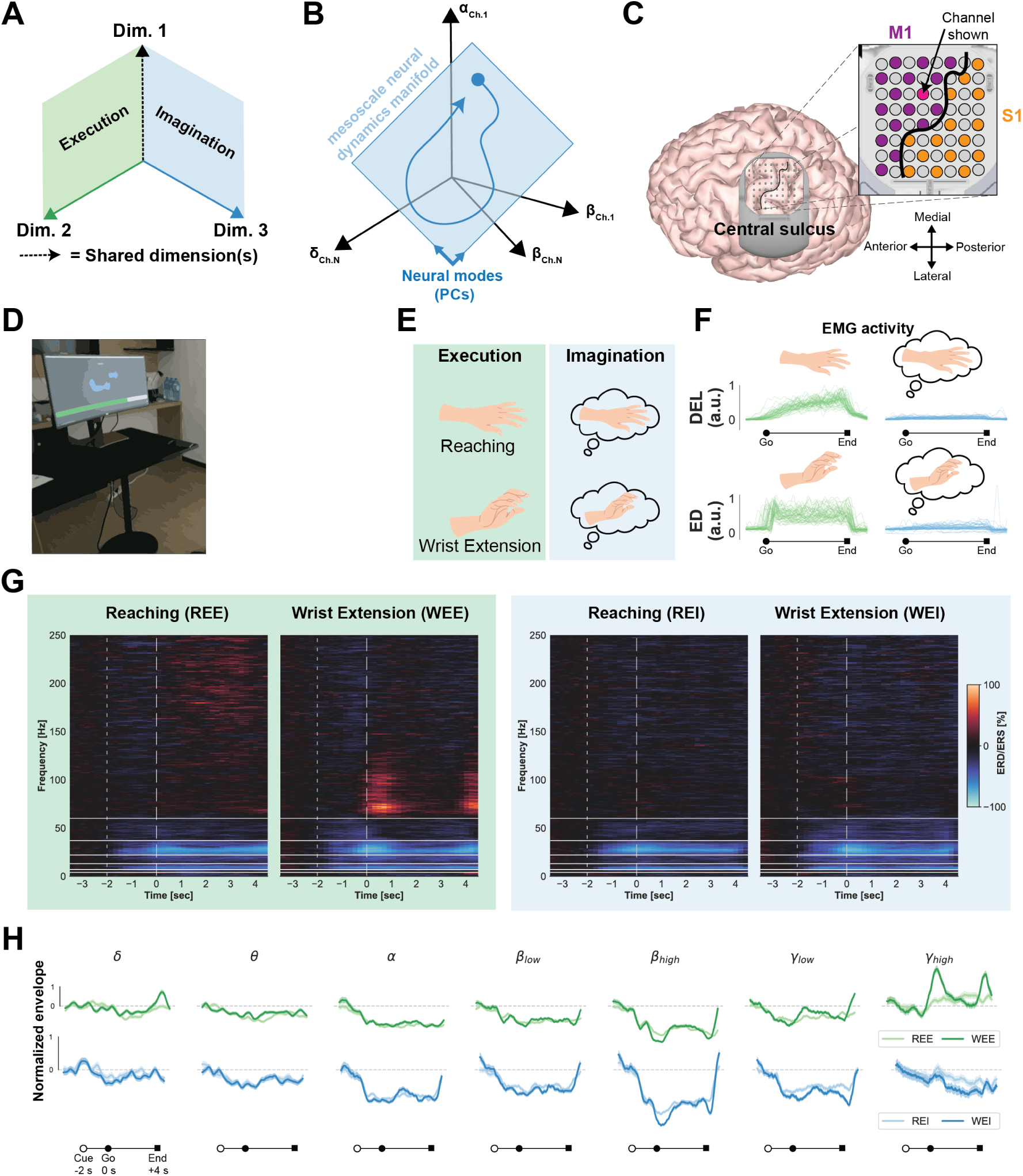
Experimental setup, paradigm and hypothesis. **A)** A sketch illustrating the hypothesis where motor imagination and execution as sharing some dimensions, while also evolving on exclusive ones. **B)** A neural manifold spanning ECoG data is defined here as a space constrained by covariance patterns among channel and frequency bands. **C)** The WIMAGINE device located on the left hemisphere over M1 and S1. **D)** The experimental setup consisted of the participant in front of a screen displaying the instructions. A progression bar helped the participant timing the movement during the trial. (Participant removed from the panel in the preprint version of the manuscript.) **E)** The paradigm consisted of two right arm movements executed, a reaching movement and a wrist extension movement, and their imaginary counterparts. **F)** EMG activity from an example session of the right lateral deltoid (DEL, to monitor activity when executing the reaching movement) and of the right extensor digitorum (ED, to monitor activity when executing the wrist extension) for executed (green) and imagined (blue) trials. **G)** Example spectrograms illustrating neural activity for an example channel (indicated in **C**) on M1 during executed and imagined movements. **H)** Normalized enveloped for all frequency bands for the same channel as in **G**. The thick lines represent the average across trials, while the shaded area represents the standard error of the mean. REE = reaching executed, WEE = wrist extension executed, REI = reaching imagined, WEI = wrist extension imagined.

A 34-year-old male with tetraplegia due to an incomplete C6 spinal cord injury was implanted with the chronic wireless ECoG WIMAGINE device [30] 14 years post-injury over the primary motor (M1) and sensory cortices (S1) (see “Participant” in Methods). The reconstructed location of the implant is shown in **Fig.1C**. During the experimental sessions, the participant sat in front of a screen displaying movement instructions alongside a progressing bar to help pace the movements (**Fig.1D**). The participant was instructed to perform both execution and imagination trials of reaching and wrist extension movements (REE = reaching executed, WEE = wrist extension executed, REI = reaching imagined, WEI = wrist extension imagined, **Fig.1E**) with his right arm, which were feasible due to the residual movement abilities of the participant (ASIA score of 5 in the elbow flexors, ASIA score of 3 on the wrist extensors). During the imagination trials, the participant was instructed to remain as still as possible. The right extensor digitorum and the lateral deltoid were chosen as target muscles for the wrist extension and reaching movements, respectively, due to the absence of EMG activity during the opposite action (**Fig. S1**). Significant muscle activity was observed during execution trials, whereas no major activity was detected during imagination trials (**Fig. 1F**).

Neural activity was modulated during both execution and imagination trials. The time-frequency decomposition for an example channel (highlighted in **Fig.1C**) is shown in **Fig.1G**. Notably, a pronounced desynchronization in the high-beta band (around 30 Hz) is observed across all conditions. To identify the neural spaces and perform all subsequent analyses, we divided the spectrum into seven frequency bands (*δ* : 1–4 Hz, *θ* : 4–7 Hz, *α* :7–13 Hz, low *β* : 13–22 Hz, high *β* : 22–37 Hz, low *γ* : 37–60 Hz, and high *γ* : 60–250 Hz) and extracted their envelopes. **Fig.1H** displays the average time-evolving envelope across trials for all frequency bands, corresponding to the same example channel.

### Execution and imagination dynamics show partial overlap at the mesoscale level

As an initial step, we performed dimensionality reduction on the original 224-dimensional space (32 channels × 7 frequency bands) to identify a global latent space that captures nearly all the variance of both execution and imagination data. To achieve this, we applied principal component analysis (PCA) separately to the trial-averaged execution and imagination data, retaining a number of principal components (PCs) that respectively explained 99% of the variance of the two datasets. We then constructed an orthonormal basis spanning the concatenation of the two transformation matrices, resulting in a 37-dimensional common latent space that captured both execution and imagination activities. To assess the alignment between these two activity types, we conducted a second PCA on the trial-averaged execution and imagination data projected onto this common space.

**Fig. 2A, B** illustrate the variance explained by each PC when PCA is applied separately to execution and imagination activities. The first few PCs in both conditions capture a significant portion of the variance of their respective conditions for both execution and imagination. However, the later PCs, which explain little to no variance in the condition they were computed on, still account for variance in the opposite condition. This trend is illustrated in **Fig. 2C**, which shows how cumulative variance is distributed within and across neural spaces. Within the same space (*Ex in Ex* and *Im in Im*), a small number of PCs account for nearly all the variance. In particular, 11 PCs were needed to explain 99% of execution variance, while 18 PCs were required for imagination. The higher number of PCs for imagination likely reflects the greater complexity of motor imagery processes. However, when projected onto the other neural space (*Ex in Im* and *Im in Ex*), variance is more evenly distributed across all components, resulting in a more gradual accumulation.

**Figure 2:**
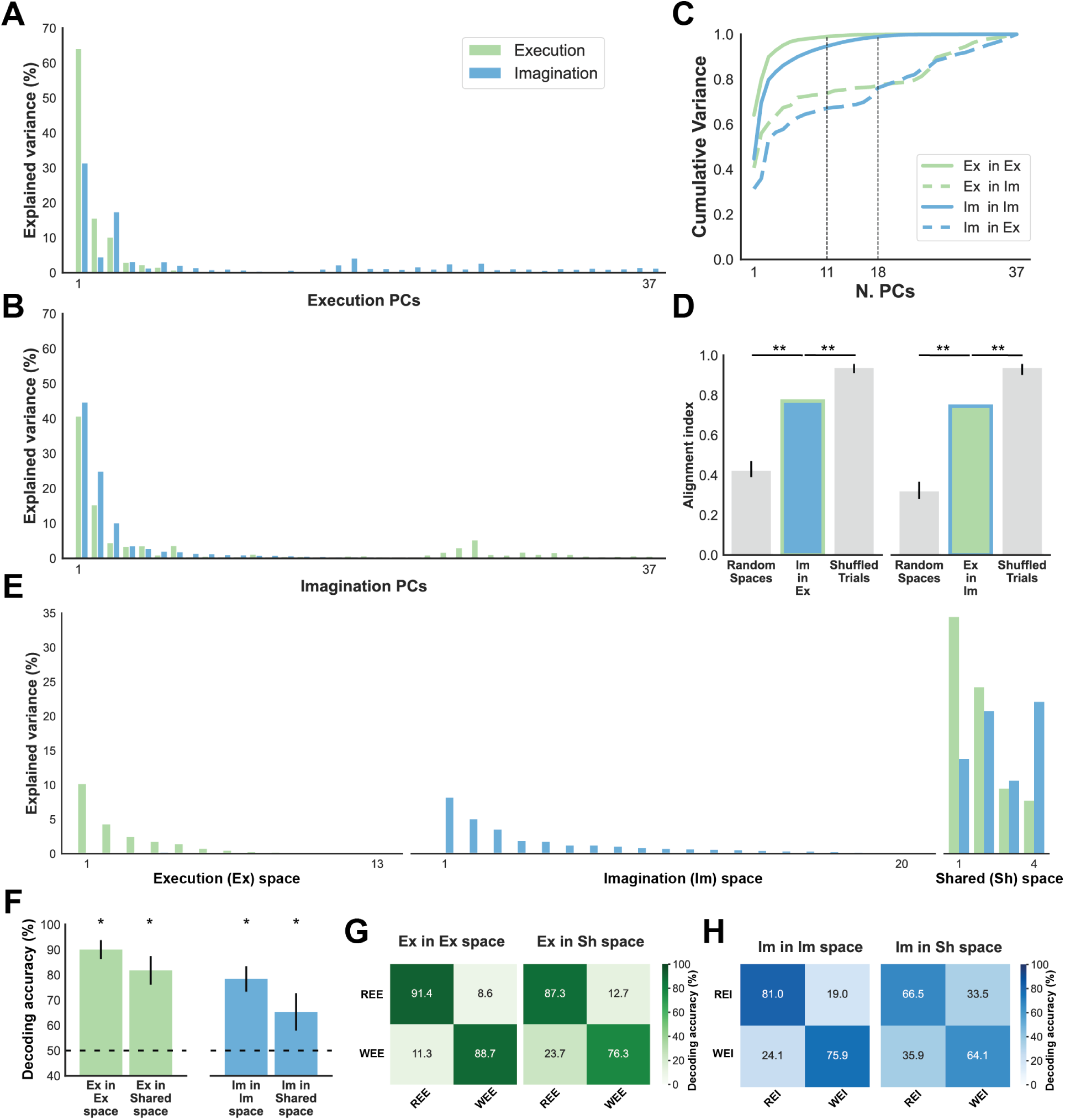
Motor execution and imagination share a neural space while also evolving onto exclusive neural spaces. **A)** Variance explained by every principal component (PC) for execution and imagination activity when performing PCA on the execution data projected onto the common latent space. **B)** Same as A but when projecting onto the PCs computed on imagination activity. **C)** Progressive cumulative variance across PCs for withing vs across spaces. **D)** Alignment index computed for both across-space projections together with a random control aiming at measuring possible alignment by chance and a shuffled-labels control aiming at evaluating alignment accounted for by trial-inherent variance. Both controls are performed with 1000 permutations and errorbars represent 0.5% and 99.5% percentiles. **E)** Decomposition of the latent space in three different space capturing only execution activity, only imagination activity, or both. **F)** Decoding accuracies when projecting single trials on the neural spaces identified. Confusion matrices for Execution **G)** and Imagination **H)**. * indicates significance at *p <* 0.01, and ** indicates significance at *p <* 0.001. REE = reaching executed, WEE = wrist extension executed, REI = reaching imagined, WEI = wrist extension imagined.

To quantify execution-imagination alignment, we used the alignment index proposed in [10, 11, 14], which ranges from 0 (total orthogonality) to 1 (full alignment). This index measures the ratio over a given number of components of variance explained across spaces (such as *Im in Ex*) relative to the variance explained within the same space (such as *Im in Im*). Using 11 and 18 dimensions for execution and imagination, respectively, we found alignment indices of 0.77 for imagination on execution and 0.74 for execution on imagination (**Fig. 2D**). This means that the first 18 PCs computed from execution activity still explain approximately 77% of the variance in imagination activity. Viceversa, using the first 11 PCs computed from imagination explain approximately 74% of execution variance.

To evaluate whether this alignment exceeded random expectations, we conducted two control analyses: (1) estimating alignment for randomly sampled subspaces constrained to match the covariance structure of the original data (yielding a distribution of random alignment indices), and (2) computing alignment after shuffling execution and imagination labels, thereby accounting for trial-to-trial variability alone and estimating how aligned the resulting spaces would be under a null model of no task structure. In both execution and imagination, our observed alignment indices were significantly higher than chance (*p <* 0.001, *N* = 1000 random samplings) but lower than the alignment obtained by shuffling labels (*p <* 0.001, *N* = 1000 permutations). These results indicate that execution and imagination activities occupy more aligned spaces than expected by chance but are not perfectly overlapping, suggesting the presence of exclusive dimensions where execution and imagination evolve separately. As a control, we repeated this analysis on intertrial activity and found complete alignment between execution and imagination, as expected (**Fig. S2**).

We then identified these shared and exclusive neural spaces by leveraging the trailing PCs of each activity and optimizing a separation process similar to [14]. To extract the exclusive execution and imagination spaces, we set a 1% explained-variance threshold to the neural trajectories of the cross-modality, respectively. This optimization yielded three orthogonal subspaces whose concatenated dimensions spanned the full latent space, providing a unified transformation that disentangled execution and imagination variance into distinct neural spaces. Specifically, we identified a 13-dimensional execution-exclusive space (hereafter referred to as the execution space) explaining 23.6% of execution variance, a 20-dimensional imagination-exclusive space (hereafter referred to as the imagination space) explaining 31.1% of imagination variance, and a 4-dimensional shared space accounting for 75.8% of execution variance and 67.2% of imagination variance (**Fig. 2E**).

Finally, we assessed the behavioral relevance of the identified neural spaces by projecting individual trials onto them and using linear discriminant analysis (LDA) to decode the two executed movements and the two imagined movements [10]. For execution data, decoding accuracy was 90.04 ± 3.75% (mean ± standard deviation across 10 cross-validation folds) when projecting onto the execution space and 81.78 ± 5.67% when projecting onto the shared space (**Fig. 2F**). For imagination data, accuracy was 78.41 ± 5.07% in the imagination space and 65.33 ± 7.39% in the shared space (**Fig. 2F**). All accuracies were significantly above chance (*p <* 0.01, permutation test, *N* = 1000 random label shuffles). The normalized confusion matrices for all classification settings are displayed in **Fig. 2G, H**.

### Frequency bands contribute differently to neural spaces

The identified execution and imagination manifolds are subspaces of the broader neural space defined by the full spectral content of the neural data. Defined by covariance patterns across channels and frequencies, these spaces let us dissect the individual contributions of each frequency band to each coding modality. To quantify individual feature contributions (i.e., channel × frequency combination) to a given neural subspace, we multiplied the transformation matrix that projects the original recordings into the global latent space by the matrix that projects the global latent space into the target subspace. To enable comparisons across spaces, we normalized the contributions within each space, obtaining the percentage contribution of each feature to that space. **Fig. S3** shows the contributions of each channel across all neural spaces and frequency bands, mapped onto the ECoG grid.

We computed the log ratios of the contributions of all features for all combinations of neural spaces (**Fig. 3A, B**). The log ratios exhibited unimodal distributions in all cases, as confirmed by the Hartigan test for unimodality [31]. This indicates that the contributions were generally homogeneously spread across features, rather than being split into distinct sets of features uniquely contributing to specific spaces, which would have resulted in bimodal distributions.

**Figure 3:**
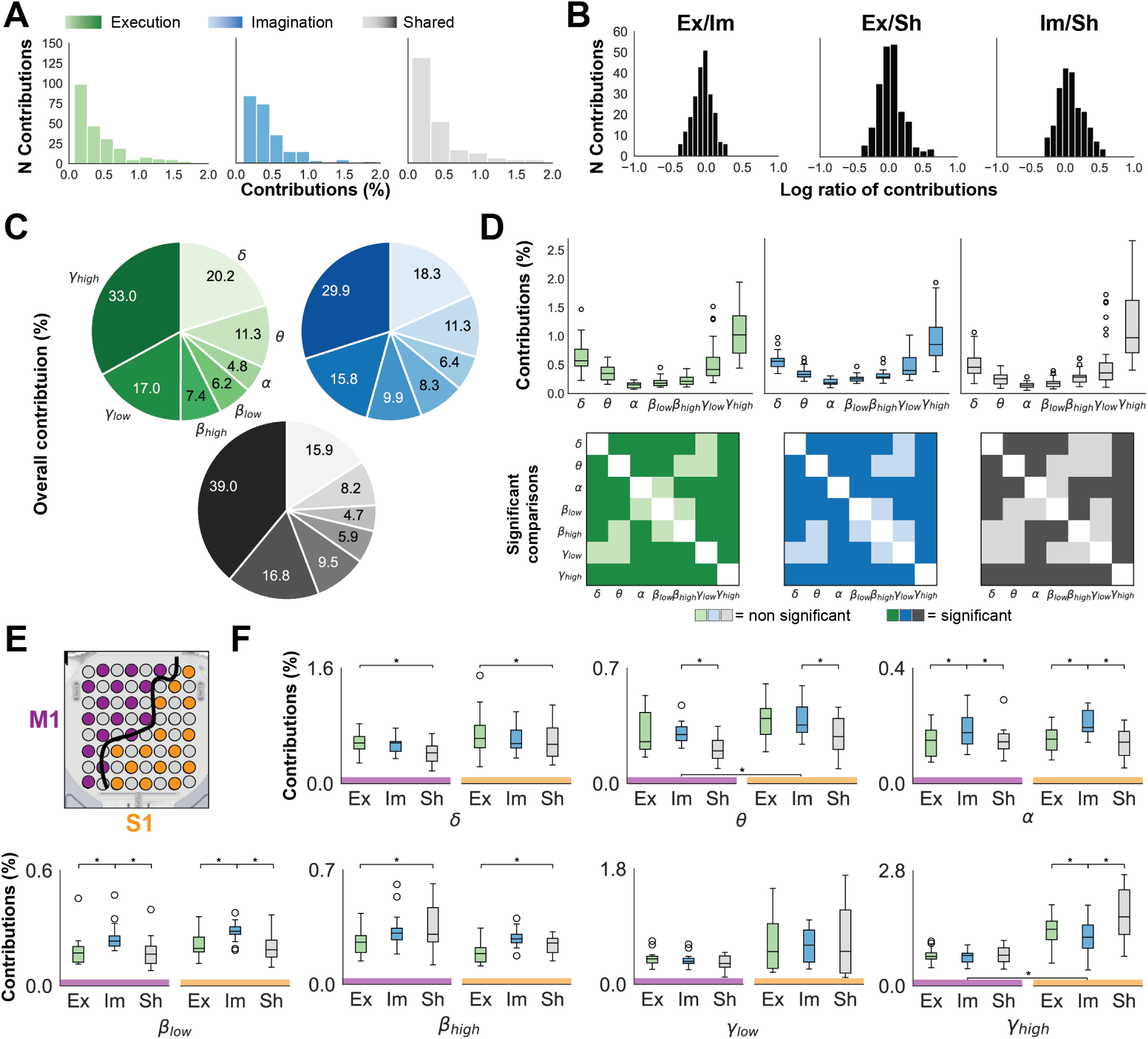
Different frequency bands contribute differently to the execution, imagination, and shared neural spaces. **A)** Distribution of contributions normalized by space. **B)** Log ratio of the three pairwise comparisons highlighting the unimodal distribution of contributions. **C)** Total variance per frequency (sum over channels). **D)** Distributions of contributions per band across channels for all neural spaces. Bottom line: Statistical significances (darker color) for the all pairwise comparisons (Repeated-Measures ANOVA, *p <* 0.01, Bonferroni corrected). **E)** Illustration of channels location over M1 and S1. **F)** Distributions of contributions across channels for all neural spaces by dividing M1 and S1 channels. * indicates significance at *p <* 0.01, mixed-effects ANOVA, Bonferroni corrected.

To summarize the overall contribution of each frequency band, we summed the contributions of all channels for each band separately (**Fig. 3C**). The largest contributions were observed in the delta (20.2% for the exclusive-execution space, 18.3% for the exclusive-imagination space, and 15.9% for the shared space) and high-gamma bands (33.0% for the exclusive-execution space, 29.9% for the exclusive-imagination space, and 39.0% for the shared space). This U-shaped pattern of frequency band contributions is clearly visible in **Fig. 3D**, which compares the distributions of contributions across channels for each space across frequency bands.

Our goal was to uncover potential patterns within specific frequency bands across different neural spaces. To achieve this, we categorized the channels based on the two brain regions covered by the implant: M1 and S1 (**Fig. 3E**). For each frequency band, we performed a mixed-effects ANOVA to test if a main effect of the cortical region, of the neural space, or of their interaction was present for a given frequency band. Then pairwise post-hoc t-tests were performed when a significant main effect was observed. The results were Bonferroni corrected for both the number of tests within each neural space and between frequency bands. We found significant statistical differences (mixed-effects ANOVA, *p <* 0.01, Bonferroni corrected) across neural spaces for all frequency bands except low gamma (**Fig. 3F**). Notably, contributions in the alpha and beta bands were significantly higher in the imagination space compared to both the execution and shared spaces, regardless of the brain region. In contrast, delta band contributions were highest in the execution space, with the shared space showing the lowest contributions. High beta activity was most prominent in the shared space. For theta, there was a regional effect, with higher contributions observed in S1 compared to M1. High gamma was the only frequency band where the interaction between neural space and region was significant, revealing statistical differences only in S1, where imagination contributions were lower than those in the execution and shared spaces.

### The exclusive imagination space remains orthogonal to execution across days

The analysis in **Fig. 2** explores the relationship between neural spaces underlying execution and imagination activity across the entire dataset. To assess the stability of this relationship and its robustness across datasets acquired over multiple days - sometimes spanning several months (**Table S1**) - we employed a leave-one-session-out (LOSO) approach. In this method, illustrated in **Fig. 4A**, neural spaces (including the global latent space and the three neural subspaces) were identified using four of the five recorded sessions (sessions 1,3,4,6, and 7, **Table S1, S2**). These identified spaces were then evaluated for their relevance to the left-out session.

**Figure 4:**
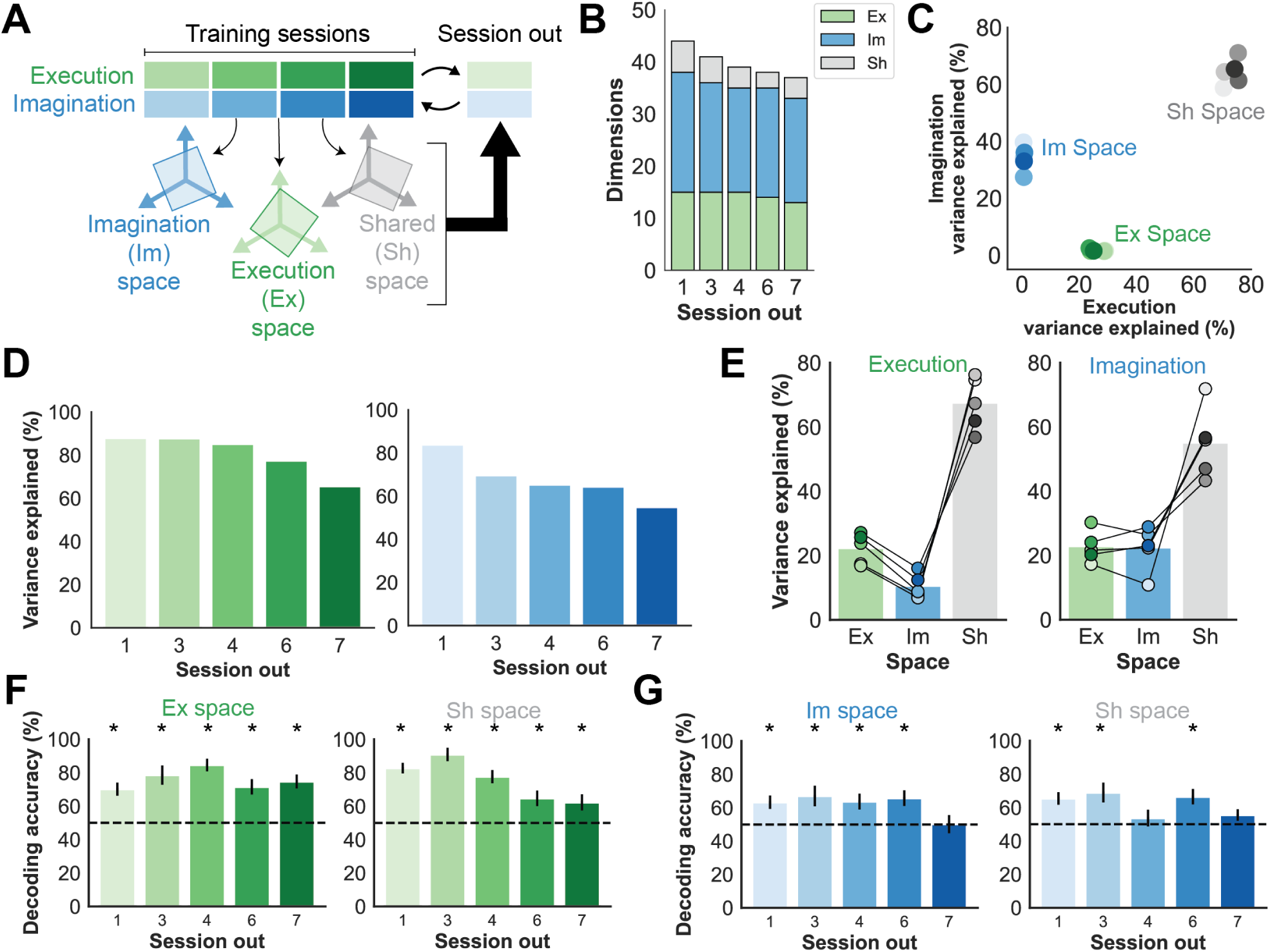
The exclusive spaces, especially the execution space, remain stable when used on new sessions of data. **A)** A sketch illustrating the approach followed in this analysis, that is a “leave one session out” approach. Neural manifolds were identified and decoders were trained on four sessions, then the session left out was used a test set. **B)** Dimensions of the three neural spaces identified for every combination of sessions. **C)** Variance explained by the three neural spaces for all combinations of sessions. **D)** Variance explained by the common latent space between execution and imagination when projecting the execution data (left) or the imagination data (right) of the session left out. **E)** Variance explained by the three neural space identified when projecting execution data (left) and imagination data (right) of the session left out onto the three neural spaces identified over the remaining sessions.**F)** Decoding of single movements when projecting the execution data onto the exclusive execution space (left) or the shared space (right). Decoders were trained on the 4 sessions and test on the session left out by bootstrapping the test set 1000 times. Error bars represent standard deviation. * indicates significance against the 99% percentile computed over the chance distribution obtained via label shuffling (1000 repetitions). **G)** Same as **F** but for pmagination data projected onto the exclusive imagination space (left) and shared space (right).

The relationship between execution and imagination activity remained stable regardless of the combination of session data tested. The dimensionality of the latent space was 40 ± 2 dimensions, demonstrating overall consistency across the tested combinations (**Fig. 4B**). Similarly, the dimensionality of the neural subspaces remained stable, with 14±1 dimensions for the exclusive execution space, 21±1 dimensions for the exclusive imagination space, and 4±1 dimensions for the shared space. Notably, the exclusive spaces consistently exhibited higher dimensionality than the shared space. The execution space accounted for 26.04 ± 2.35% of the variance in execution data, while the imagination space explained 34.07 ± 4.01% of the variance in imagination activity. The shared space explained 73.26 ± 2.28% and 64.21 ± 4.14% of execution and imagination data, respectively (**Fig. 4C**). Complementary results from a similar analysis including neural spaces identified from individual sessions, are presented in **Fig. S4**.

When evaluating how these neural spaces generalize across days, we found that the global latent space (**Fig. 4D**) explained 80.85 ± 8.50% of the variance in execution activity not used to define the space but only 67.64 ± 9.43% of the variance in imagination activity. Further projecting the neural activity from the left-out session onto the individual neural subspaces (**Fig. 4E**) revealed that the execution variance captured by the global latent space was distributed as follows: 22.18 ± 4.27% explained by the exclusive execution space, 10.41 ± 3.39% by the exclusive imagination space, and 67.41 ± 7.41% by the shared space. For imagination activity, the variance explained by the global latent space was divided into 22.72 ± 4.38% by the exclusive execution space, 22.32 ± 6.20% by the exclusive imagination space, and 54.96 ± 9.89% by the shared space. These results highlight the exclusive imagination space as a “functional null space” for execution activity, capturing minimal variance and thus being partially orthogonal to it. However, this asymmetry does not hold for the exclusive execution space, as both exclusive spaces explain similar amounts of imagination activity variance.

### Task relevance of neural spaces persists across days

We aimed to determine whether the neural subspaces could generalize in terms of behavioral relevance, specifically assessing whether the variance they captured was adequate to distinguish between the reaching and wrist extensions movements when tested on unseen data. To test this, single trials from the sessions used to identify the neural spaces were concatenated and projected onto these subspaces. We then trained a classifier on this dataset and tested it on trials from the left-out session, also projected onto the neural subspaces. A robust and stable neural representation would allow the classifier to generalize and accurately classify trials from the unseen dataset. For every LOSO combination, the classification accuracy was calculated as the average of 1000 resamplings of train and test trials.

For execution (**Fig. 4F**), projecting onto the exclusive execution space resulted in an average accuracy of 76.0 ± 5.2% across LOSO combinations, while projecting onto the shared space yielded an accuracy of 75.7 ± 10.8%. In all cases, accuracies were significantly above chance (p *<* 0.01, permutation test, *N* = 1000). For imagination (**Fig. 4G**), projecting onto the exclusive imagination space resulted in an average accuracy of 62.0 ± 6.1%, and projecting onto the shared space yielded 62.0 ± 6.2%. All combinations were significantly above chance (*p <* 0.01, permutation test, *N* = 1000) except when session 7 (for decoding onto both spaces) or session 4 (for decoding onto the shared space) were left out.

### Extending neural space identification to additional imagined movements

After identifying exclusive and shared neural spaces for execution and imagination at a mesoscale level, we next investigated whether similar spaces could be detected when introducing additional imagined movements that could not be physically executed. While this approach disrupts the symmetry of directly comparing the same movements across execution and imagination, it holds significant translational potential. For participants with residual movement abilities, identifying a neural space that selectively captures imagination-related activity, while excluding variance shared with their executable movement repertoire, could be particularly valuable for BMI applications. To investigate this, the participant was asked to imagine additional movements, specifically walking and grasping (**Fig. 5A**), during three of the five sessions dedicated to single-movement tasks (see **Table S2**). As expected, also during these imagined movements, the EMG activity of the lateral deltoid and extensor digitorum was flat (**Fig. 5B**). The time-frequency decomposition of the two added movements are presented in **Fig. 5C** for the same example channel shown in **Fig. 1H** (see **Fig. S5** for the envelopes of this example channel).

**Figure 5:**
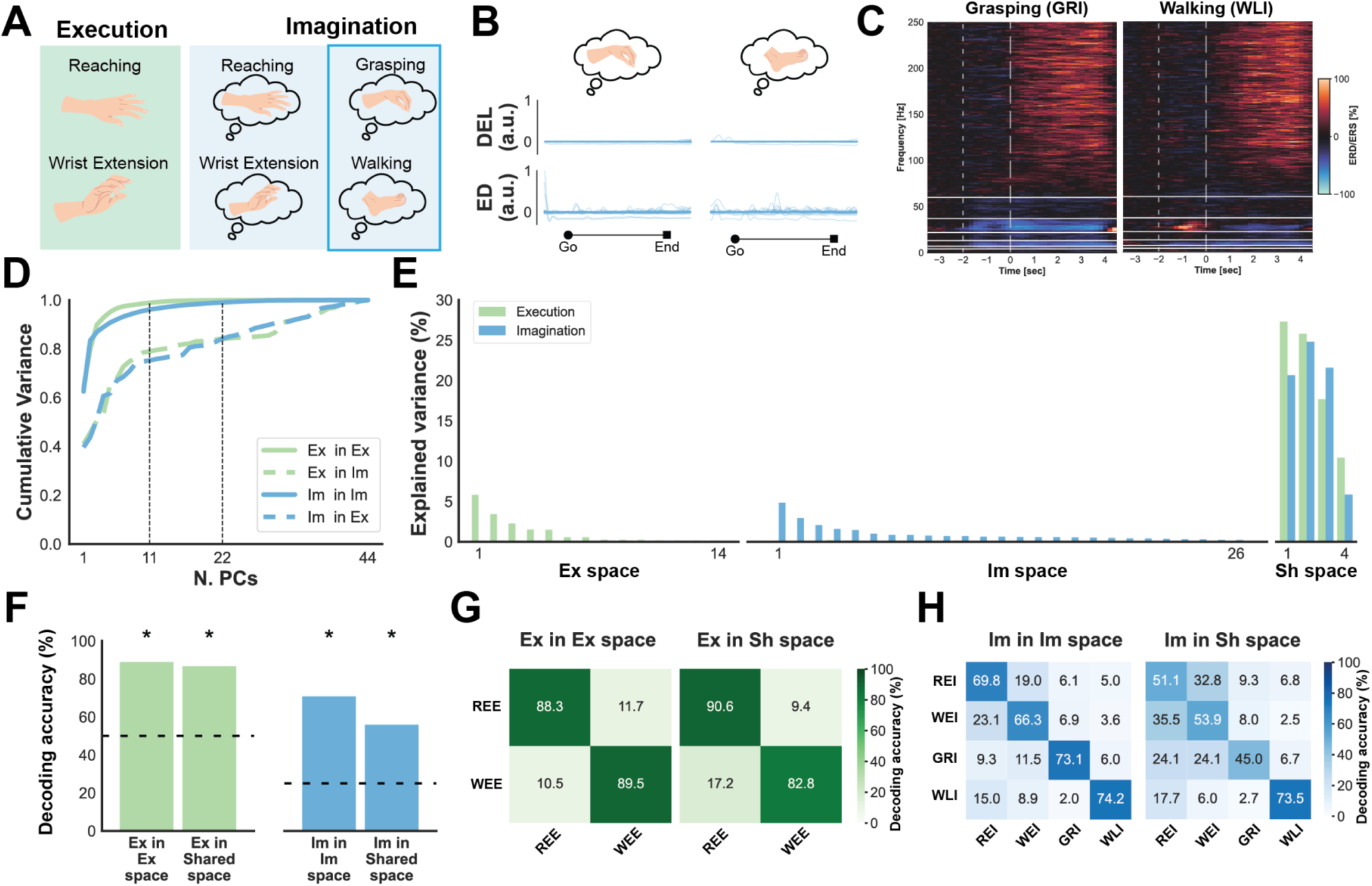
Similar spaces can be identified also when adding more imagined movements. **A)** As a second paradigm, two additional imagined movements were considered: grasping and walking. **B)** EMG activity from an example session during imagined grasping (left) and walking (right) trials for the target muscles shown in Fig. 1F. DEL: lateral deltoid, ED: extensor digitorum. **C)** Time-frequency decomposition for the same example channel as in Fig. 1H. **D)** Cumulative variance across all PCs for execution and imagination projections within and across spaces when performing PCA over the common latent space. Decomposition of the latent space in the three separate neural spaces. **F)** Decoding accuracies when projecting single trials on the neural spaces identified. * indicates significance against the 99% percentile of the chance distribution computed over 1000 label shuffles. Theoretical chance level is shown by the dashed line. Confusion matrices for execution **G)** and imagination **H)**. REE = reaching executed, WEE = wrist extension executed, REI = reaching imagined, WEI = wrist extension imagined, GRI = grasping imagined, WLI = walking imagined.

For these three sessions, we repeated the procedure described above to identify the neural spaces, this time including two executed movements and four imagined movements. **Fig. 5D** illustrates the cumulative explained variance within and across spaces when applying PCA separately to execution and imagination data, after projecting them onto a new 44-dimensional common latent space (**Fig. S6**). Similarly to the previous analysis, the trailing PCs of one space still captured a significant portion of variance from the other. In this case, 22 dimensions were required to explain 99% of the variance within the imagination space, compared to 18 in **Fig. 2C**. This slight increase in dimensionality, along with the expansion of the common latent space, reflects the inclusion of additional imagined movements, leading to greater variance within the neural data. **Fig. 5E** presents the separation of the 44 dimensions of the latent space into 14 dimensions explaining exclusively 17.3% of the execution variance, 26 dimensions explaining exclusively 25.7% of imagination variance, and 4 dimensions composing the shared space explaining 81.2% and 72.9% of execution and imagination variance, respectively.

We conducted a decoding analysis (**Fig. 5F-H**) on single trials projected onto the corresponding neural spaces to evaluate whether these newly identified spaces retained behavioral information. In this case, the theoretical chance level when classifying the imagined movements was 25%, given the presence of four classes instead of two. The analysis yielded accuracies of 88.92 ± 4.01% (mean ± standard deviation across 10 cross-validation folds) and 86.69 ± 5.99% for executed trials projected onto the exclusive-execution and shared spaces, respectively. For imagined movements, we achieved accuracies of 70.86 ± 4.58% and 55.89 ± 5.13% for the exclusive-imagination and shared spaces, respectively.

### Exclusive neural spaces enable the separation of concurrent imagination and execution activity

Once the neural spaces for both execution and imagination were identified, we hypothesized that the exclusive neural subspaces could effectively disentangle the corresponding activities during concurrent execution and imagination. To test this, the participant took part in additional dual-task sessions (see **Tables S1, S2, S3**), where he was instructed to perform a movement while simultaneously imagining another. The participant was explicitly told to execute both tasks concurrently rather than sequentially.

Within the full experimental paradigm, several execution-and-imagination pairings were possible. In particular, we compared the encoding of different imagined actions while subjects performed the same physical action, and the encoding of different executed actions while they imagined the same action. There were three possible combinations involving the execution of a reaching movement: reaching while imagining a wrist extension, imagining grasping, or imagining walking (**Fig. 6A**, point 1). Two combinations were considered when performing the wrist extension movement: extension while imagining reaching or imagining walking. The combination of executing wrist extension while imagining grasping was excluded, as it was deemed too counterintuitive and unnatural (**Fig. 6A**, point 2). Among these combinations, we also focused on the comparison between imagining walking while executing a reaching movement or a wrist extension (**Fig. 6A**, point 3).

**Figure 6:**
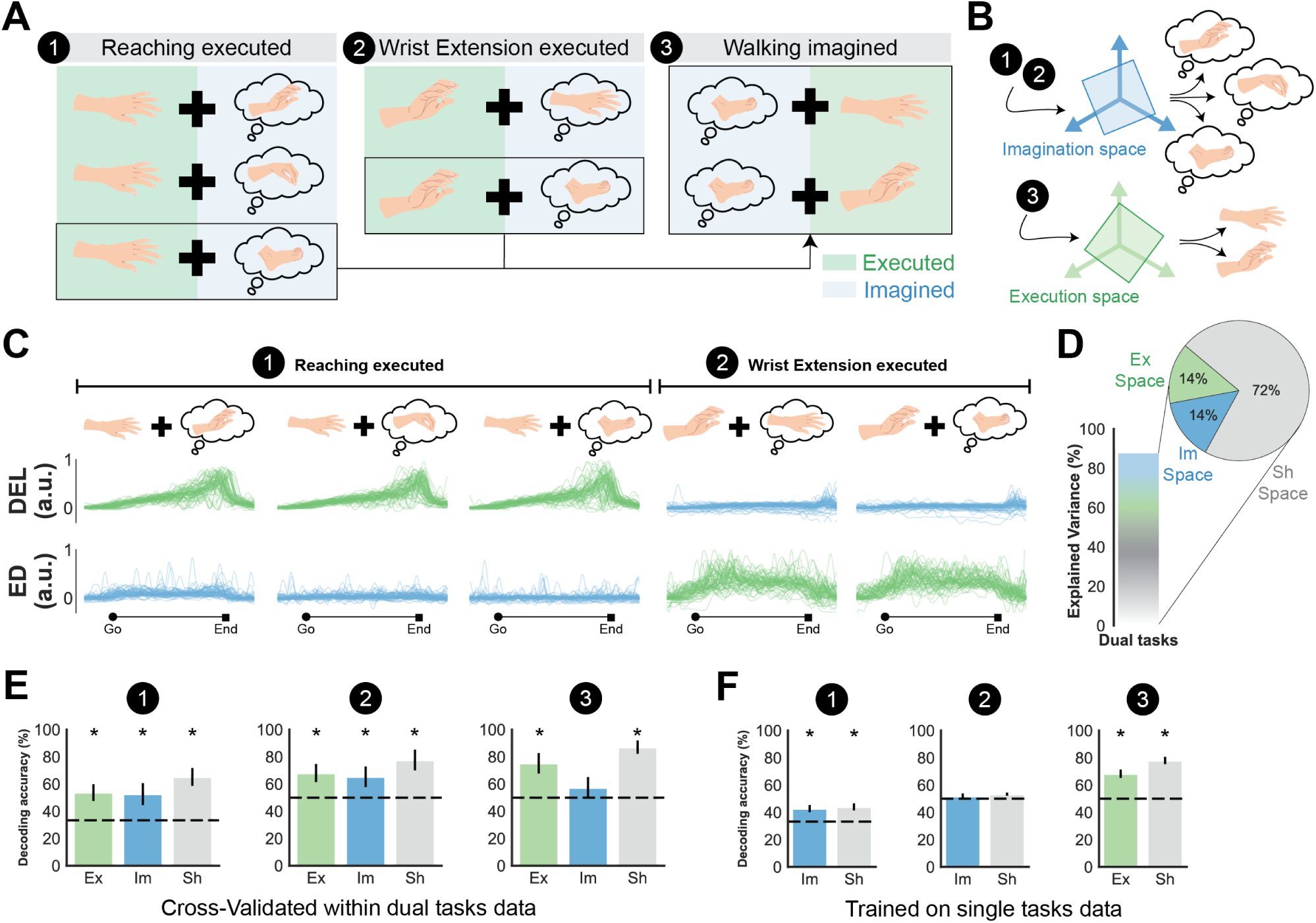
The exclusive neural spaces can help disentangle execution and imagination activity when occurring together. **A)** Experimental paradigm concerning the dual tasks approach. Note that there was no dual task involving the execution of the wrist extension while imagining grasping because considered counterintuitive. **B)** Sketch illustrating our hypothesis seeing the exclusive spaces as a potential tool to help disentangle executory and imaginary components when dual-tasking. **C)** Recorded EMG activity of the lateral deltoid (DEL) and extensor digitorum (ED) during an example session of dual tasks. The muscle targeting the executed movement is shown in green (DEL for reaching and ED for wrist extension movements), while the non-target muscle of each executed movement is presented in blue. **D)** Explained variance by the latent space found on the single tasks from Fig. 5 (bar plot) and subdivision of the explained variance into the three different neural spaces (pie chart). **E)** Decoding accuracies obtained when projecting the dual task data onto the different neural spaces, and then cross-validating within the dual task data (i.e. the classifier can learn from the dual task distribution). The dashed line represents the theoretical chance level. **F)** Decoding accuracies obtained when projecting the dual task data onto the different neural spaces, training a classifier onto the single tasks and testing it (by bootstrapping) on the dual task data. The dashed line represents the theoretical chance level. * indicates significance against the 99% percentile of the chance distribution found by labels shuffling.

In cases 1 and 2, where the executed movements remain constant and the imagined movements change, we expect to distinguish the imagined movements by projecting the neural data onto the exclusive imagination space. Conversely, in case 3, where the executed movements change while the imagined movement remains constant, we expect that the exclusive execution space will enable us to distinguish between the two executed movements. **Fig. 6B** illustrates our hypothesis.

Examples of neural activity across all frequency bands and movement combinations are shown in **Fig. S7**. EMG activity of the target muscles highlighted the strong modulation of the muscle engaged in executed movements and the lack of activity in the other muscle (**Fig. 6C**). When projecting the trial-averaged neural activity from dual-task trials onto the neural subspaces previously identified from the single-task datasets, we found that the common latent space explained 89% of the variance. Of this, 14% was explained by the exclusive execution and imagination spaces each, while 72% was explained by the shared space (**Fig. 6D**).

To assess whether different movements could be decoded, we employed two distinct approaches. In both cases, dual-task neural data were projected onto neural spaces derived from the single-task dataset. The first approach focused on evaluating the reliability of the neural representations themselves by performing cross-validation within the projected dual-task data, allowing classifiers to adapt to the new data distribution. **Fig. 6E** presents the decoding accuracies for this approach. In both cases 1 and 2, the imagined movements (a three-class problem for case 1, and binary classification for case 2) could be decoded significantly above chance (*p <* 0.01, permutation test, *N* = 1000) when projecting onto all spaces. Specifically, for case 1 (reaching executed), we achieved 53.7 ± 6.1% (mean ± standard deviation across 10 folds) accuracy when projecting onto the exclusive execution space, 52.6 ± 8.0% when projecting onto the exclusive imagination space, and 65.2 ± 6.6% when projecting onto the shared space. For case 2 (executed wrist extension), the accuracies were 68.0 ± 6.6% for the exclusive execution space, 65.3 ± 7.6% for the exclusive imagination space, and 77.6 ± 7.6% for the shared space. Notably, decoding was possible even when projecting onto the exclusive execution space. In contrast, for case 3 (walking imagined), the decoding accuracy of the executed movements was significantly higher than chance (*p <* 0.01, permutation test, *N* = 1000) only when projecting onto the exclusive execution space (75.2 ± 7.5%) and the shared space (87.1 ± 4.8%), but not when projecting onto the exclusive imagination space (57.3 ± 7.8%), consistent with our hypothesis.

In the second decoding approach, we tested whether a classifier trained on the single-task data could generalize to the dual-task condition by detecting shared neural dynamics. Specifically, a classifier pre-trained on the single-task dataset was applied directly to the projected dual-task data. Building on the first approach, this method moves towards the concept of future “plug and play” BMIs, where neural spaces could be identified and decoders could be trained during initial calibration sessions, and subsequently applied across sessions without retraining. **Fig. 6F** shows the decoding results for this approach. Note that for cases 1 and 2, the results for projections onto the exclusive execution space, and for case 3, the results for projections onto the exclusive imagination space, are not reported, as these would correspond to random classifiers by construction. This is because they would be trained on the same data used to identify the neural spaces, which explains only about 1% of the variance.

Our analysis revealed that decoding was significantly above chance (p *<* 0.01, permutation test, N = 1000) for all conditions except case 2 (wrist extension executed). This indicates that a classifier trained on single-task data to distinguish imagined walking and grasping was unable to differentiate these movements in the dual-task setting. In case 1 (reaching executed), accuracy reached 42.8 ± 2.6% (mean ± standard deviation across 1000 resamplings) when projecting onto the exclusive imagination space, and 44.1 ± 2.5% when projecting onto the shared space. For case 3 (walking imagined), accuracy was 68.3 ± 3.0% when projecting onto the exclusive execution space and 78.1 ± 2.6% when projecting onto the shared space. Notably, these results remained consistent regardless of the duration of time windows or the amount of variance retained during decoding (**Table S4**).

## Discussion

In the context of BMIs, a key challenge lies in disentangling the neural activities underlying motor imagination and execution, particularly when they occur simultaneously, as this overlap can lead to neural cross-talk and compromise the independent control of external devices. Addressing this challenge has important implications in improving BMI control in individuals with residual motor function. To tackle this, we turned to mesoscale signals and adapted the neural manifold framework, originally developed for spiking activity, to ECoG recordings. This allowed us to validate and extend prior findings on the structure of motor representations, and to demonstrate the suitability of mesoscale neural dynamics for coordinated and independent control in BMI applications.

When transitioning from spiking activity to mesoscale activity, it remains unclear whether neural representations identified at higher definition generalize to the mesoscale level when recording from broader areas and at a lower spatial resolution. For instance, differences were observed when considering a preserved manifold encoding a repertoire of movements, where temporal dynamics were maintained in spiking activity [8, 32] but not necessarily in ECoG recordings [33]. Our findings succeed in extending a manifold approach to mesoscale neural dynamics measured via ECoG, providing insights into the relationship between the encoding of motor execution and imagination. In agreement with studies on spiking activity that explore motor execution and its relationship to motor imagination [14], and observation [10], we also identify a subspace of shared covariance between execution and imagination (**Fig. 2D**). We found that the shared neural space has significantly lower dimensionality than the exclusive spaces (**Fig. 2E**), suggesting that the exclusive spaces capture more intricate patterns and dynamics, as previously suggested [14]. Decoding analyses (**Fig. 2F-H**) confirmed that these subspaces retain behaviorally relevant information, emphasizing their functional significance. It is worth noting that our decoding efforts aimed primarily to demonstrate the presence of task-related information, and further refinement of models could potentially improve decoding performance within these neural spaces.

The adaptation of this framework to mesoscale neural dynamics also involves defining neural spaces as covariance patterns across channels and frequency bands. We observed a U-shaped pattern of frequency contributions (**Fig. 3C-D**), with delta and high-gamma bands showing the highest contributions. This is consistent with previous studies that highlight the role of these frequency bands in motor control [22, 33] and their correlations with underlying neuronal population dynamics [34–38]. When comparing the frequency band contributions across spaces (**Fig. 3F**), we found that the imaginary space exhibited significantly higher contributions in the alpha and low-beta bands. These findings align with studies that have identified modulations in these bands for motor imagery decoding in BMIs [1, 39, 40]. We also examined contributions from the M1 and S1 regions. Notably, there was little difference between their contributions, supporting the idea of a role for the primary somatosensory cortex in cognitive imagery and motor production without sensation [41]. A significant interaction between neural spaces and brain regions was observed for high gamma activity, reinforcing its key role in motor modulation [1, 33]. In this case, the shared and exclusive execution spaces showed higher contributions than the exclusive imagination space only for the channels located over S1, which might highlight the absence of sensory feedback during the imaginary process. Interestingly, from the distribution of contributions on the ECoG grid (**Fig. S3**), we observed that the contributions of the high beta band to the shared space appear to be higher in M1 than in S1, likely reflecting the strong modulation observed across conditions in this frequency band. Similarly, a localized increase in the contribution from the more lateral channels in S1 appears for the low-gamma band, which may be associated with a more localized modulation of the hand and arm regions within the somatotopic organization of the sensorimotor cortex. In this study, we chose to analyze the combined regions to maintain a higher dimensionality in the original recording space and to incorporate potential covariance patterns between regions that may be essential for identifying neural spaces, but we acknowledge that an alternative approach could be to perform the whole analysis onto the two regions separately.

A key focus of our analysis was assessing the stability and robustness of neural spaces across different sessions recorded in different days (**Fig. 4A**). The partial overlap between the neural representations of motor execution and motor imagery remained consistent across session combinations (**Fig. 4B-C**) and within individual sessions (**Fig. S4**). However, when applied to data recorded on different days, neural spaces faced challenges in consistently capturing variance, highlighting the complexity of generalizing these representations over time. There are two essential properties of these neural spaces: they are orthogonal to each other and representative of their corresponding activity, capturing task-relevant variance. These properties are independent. For example, the exclusive imagination space remained partially orthogonal to execution activity, capturing minimal execution-related variance (**Fig. 4E**). In contrast, the exclusive execution space failed to maintain strict orthogonality, capturing a significant portion of imagination-related variance. This may be due to the inherently higher variability of motor imagery compared to execution, which tends to be more stable across sessions. Despite this, both exclusive spaces demonstrated strong decoding performance across sessions, indicating that task-relevant information was well-preserved. Their performance was comparable to, or even surpassed, that of the shared space, despite the latter explaining a larger proportion of variance. Furthermore, the exclusive spaces yielded more consistent results on across-session decoding. Notably, for execution, decoding performance within the shared space exhibited greater fluctuations, whereas for imagination, the shared space failed to achieve significant decoding in one additional session combination, a limitation not observed in the exclusive space. While we recognize that greater stability might have been achieved by applying session alignment before projecting neural subspaces, we opted for a simpler approach tailored to translational applications [14, 27, 42]. The goal of this approach is to mimic the process of identifying neural spaces in calibration sessions, allowing them to be directly applied in future BMI sessions. Moreover, recent evidence suggests that accumulating data across multiple sessions could help establish a more stable, time-independent neural representation. This, in turn, may facilitate the development of generalizable classification boundaries that remain consistent over time [43].

Building on the translational perspective, we tested the proposed approach in scenarios where there is an imbalance between the number of executed and imagined movements (**Fig. 5**). This is particularly relevant for applications involving patients with residual motor capabilities, where the goal is to isolate neural variance associated with executed movements while maximizing the decoding of imagined movements. Our results demonstrated that task-relevant neural spaces could still be identified in such cases. Notably, the dimensionality of the exclusive execution and shared spaces remained stable, while that of of the exclusive imagination space increased, as expected, with the inclusion of two additional imagined movements. Interestingly, the decoding analysis (**Fig. 5H**) revealed that the two additional imagined movements, grasping and walking, were the best classified when trials were projected onto the exclusive imagination space, likely reflecting their stronger neural modulation as shown by the time-frequency decompositions (**Fig. 5C**). This stronger modulation may be attributed to the fact that these two movements were part of the participant’s BMI training protocol. Another possible and complementary explanation is that these two movements may fall into the category of attempted movements, rather than being purely imagined. This would imply that the participant was actively trying to perform the movements, which ultimately failed due to the spinal lesion. Conceptually, an attempted movement lies between executed and imagined movements: it lacks the sensory feedback component typical of executed actions, yet it also lacks the inhibitory component preventing overt movements characteristic of motor imagery [5]. From an application standpoint, however, attempted movements should be grouped with imagined movements in the context of future BMIs for motor restoration developed under our framework, as both represent “mental tasks” to be decoded concurrently with goal-directed executed movements. In this sense, regardless of whether a task is attempted or imagined, our aim would be to identify a neural subspace that exclusively covers the activity related to these “mental tasks”, thus disentangling it from that of actual motor execution. We also observed that when projected onto the shared space, imagined walking showed the highest decoding accuracy, whereas imagined grasping had lower accuracy. Since the shared space inherently incorporates variance linked to executed movements, we speculate that this difference arises because the shared space predominantly captures neural activity associated with upper limb movements, making imagined walking more distinct compared to the other movements.

In **Fig. 6**, we showed the potential of condition-specific neural spaces as effective tools for separating execution and imagination activities when they occur simultaneously. As mentioned above, this becomes directly relevant in scenarios where a participant with residual movements engages in a BMI task. The objective is to decode imagined movements while allowing concurrent execution of other movements, ultimately improving motor control and enhancing the overall BMI experience. These findings, presented in **Fig. 6E**, align with the stability analysis results shown in **Fig. 4**. Given the challenge of maintaining orthogonality to imagination activity when projecting new data onto the execution space, we observed significant decoding performance even when projecting trials onto the exclusive execution space in cases 1 and 2 (reaching executed and wrist extension executed). This result emerged when the classifier was allowed to adapt to the new data distribution of dual tasks. It is important to note that this effect might be further influenced by potential variations in the execution of the same movement while imagining different movements. On the other hand, the exclusive imagination space remains stable and functionally orthogonal, successfully preventing the decoding of the two executed movements when projected onto this space (case 3). Moreover, it is noteworthy that the decoding performances of the exclusive spaces were comparable to those of the shared space, despite the shared space capturing more variance. This suggests that the exclusive spaces are more task-relevant than the shared space, a trend that aligns with other decoding results presented in this study (**Fig. 4F-G**, **Fig. 5F-H**). Crucially, the results in **Fig. 6F** reveal that the neural patterns identified by the classifiers in the single-task datasets can also be detected and utilized when applied to dual-task data. Although no successful decoding was achieved for wrist extension execution (case 2), we observed low but consistently significant decoding performance in cases 1 and 3. This further emphasizes the task-relevant nature of the identified neural spaces and their potential in disentangling concurrent neural activities.

This study demonstrates that a neural manifold approach can be adapted to mesoscale neural activity, enabling the identification of spaces related exclusively to motor execution and motor imagination. While the framework shows promise for improving BMIs, challenges remain in achieving robust generalization across sessions, and further validation with additional patients is needed. The influence of electrode placement on cortico-cortical covariance patterns is also not fully understood, and variations may yield different geometric properties. A key next step will be to evaluate the feasibility of this approach in real-time settings, where decoding would rely on shorter, ongoing time windows rather than full-movement trajectories. In this context, incorporating closed-loop control with neurofeedback may enhance classifier performance. By reinforcing voluntary modulation within the exclusive imagination space, neurofeedback may improve the separability of imagined movement representations and increase the reliability of BMI control [1, 22, 43–45]. Looking ahead, it will be important to assess how this framework compares with alternative approaches, such as complex decoders or artificial neural networks, trained and tested on single or dual task datasets. These methods may also capture implicit manifolds for execution and imagination. Investigating their relative generalization performance as the repertoire of imagined and executed movements expands will be key to understanding the strengths and limits of each approach.

This novel framework opens up new possibilities for clinical and neuroengineering applications. By disentangling imagined activity from concurrent execution, it could support mobility and independence in paralyzed patients with residual motor function. In particular, our study introduces a geometric method to identify an execution-null space, which could be exploited to control additional degrees of freedom during concurrent movements, thus marking a promising step toward more versatile and effective motor restoration strategies.

## Methods

### Participant

All data in this study were recorded from a 34-year-old male participant with incomplete tetraplegia resulting from a C6 spinal cord injury. Fourteen years post-injury, the participant was bilaterally implanted in November 2019 with the WIMAGINE epidural chronic wireless ECoG device as part of clinical trial NCT02550522 [46]. The participant retained residual movements, particularly in the elbow flexors (American Spinal Injury Association Scale - ASIA score of 5) and wrist extensors (ASIA score of 3) [47]. This study was carried out 5 years after implantation.

The WIMAGINE device consists of 64 planar electrodes, each 2.3 mm in diameter, with an inter-electrode spacing of 4–4.5 mm [30]. Due to limited data rates from the restricted radio link and a malfunction of the right implant, 32 electrodes from the left implant were selected in a checkerboard-like pattern for the experimental sessions of the present study. Since the implantation, the participant has been trained to control various real and virtual effectors via a brain-machine interface (BMI), utilizing mental imagery and his residual movements.

### Experimental setup

The dataset for this study includes recordings from eight experimental sessions conducted over approximately 18 months (**Table S1**). Trials in these sessions involved either executed or imagined movements with the right arm. The sessions were divided into two types: single-tasks, which featured only one movement (either executed or imagined), and dual-tasks, which combined one executed movement with one imagined movement. Specifically, sessions 1, 3, 4, 6, and 7 were single-task sessions, while sessions 2, 5, and 8 were dual-task sessions (**Table S2**).

During the sessions, the participant sat in front of a screen that displayed instructions for each trial. A trial began with a pictogram representing the movement for that trial, initially shown in gray as a Cue signal. After 2 seconds, the pictogram turned bright blue, signaling the Go phase. The participant then executed or imagined the movement for 4 seconds, aided by a progression bar displayed below the pictogram to assist with timing. If the movement was completed before the 4 seconds elapsed, the participant was instructed to hold the final position, both in execution and imagery tasks. Each trial concluded with a 4-second intertrial interval, resulting in a total trial duration of 10 seconds.

The movements used in the experiment included reaching and wrist extension of the right arm, contralateral to the ECoG implant, which were selected based on the participant’s residual motor capabilities. For imagined movements, the tasks additionally included imagined walking and imagined grasping, which were introduced starting from session 4. Beginning with this session, EMG activity was recorded to monitor the participant’s compliance with the tasks. The EMG signals were collected in bipolar fashion using the Noraxon Delsys system connected to a LabJack data acquisition device, with six channels over the flexor digitorum, extensor digitorum, triceps, biceps, trapezius, and lateral deltoid muscles. The extensor digitorum and lateral deltoid were selected as the target muscles to assess contraction levels during wrist extension and reaching movements, respectively.

Each session was organized into blocks, which varied depending on the type of session. In single-task sessions, three types of blocks were included: one containing executed and imagined trials for reaching movements, another containing executed and imagined trials for wrist extension movements, and a third with only imagined trials for walking and grasping movements. Each block contained either 20 or 30 trials, balanced across conditions and presented in random order.

In dual-task sessions, two types of blocks were used. In the first, executed reaching movements were paired with one of three imagined movements: wrist extension, walking, or grasping. In the second, executed wrist extension movements were paired with either imagined reaching or imagined walking. Trials combining executed wrist extension with imagined grasping were excluded due to the task difficulty. Within each block, every condition was repeated either 10 or 12 times in random order. The order of the blocks was randomized for each session.

### Data acquisition and signal preprocessing

Data were acquired using the WIMAGINE implant [30]. Preprocessing was initially performed by the implant, which applied analog bandpass filtering with a bandwidth of 0.5–300 Hz. After digitization, a digital low-pass FIR filter with a cutoff frequency of 292.8 Hz was applied to further refine the signals.

Subsequently, the ECoG data were processed offline following a similar approach to that described in [33]. First, signals were bandpass filtered using a 5th-order Butterworth filter with cutoff frequencies of 1–250 Hz. A common median reference filter was then applied across channels to mitigate the effects of shared noise or artifacts. Next, the envelope of the signal was extracted for seven frequency bands spanning the full signal range: *δ* (1–4 Hz), *θ* (4–7 Hz), *α* (7–13 Hz), low *β* (13–22 Hz), high *β* (22–37 Hz), low *γ* (37–60 Hz), and high *γ* (60–250 Hz). This was achieved by bandpass filtering the signals within the corresponding frequency ranges using a 5th-order Butterworth filter, followed by computing the magnitude of the Hilbert-transformed signal. To reduce noise, particularly in the higher frequency bands, the envelope was smoothed using a 500-millisecond moving mean filter. Data were then subsampled to 100 Hz and mean centered per block.

Trials were then epoched by aligning the signals to the Go signal. For this study, the entire movement period, lasting 4 seconds, was analyzed. To facilitate the main analysis, trials were concatenated across sessions, creating a single dataset. Trials exhibiting artifacts, characterized by sudden signal jumps, were visually inspected, identified and excluded using a first-derivative threshold-based approach (derivative ≥ 0.3). More details on the total number of trials analyzed per experimental condition and session are provided in **Table S3**.

For normalization, signals from each session, channel, and frequency band were z-scored relative to baseline activity to remove session-dependent fluctuations. The baseline was defined as the signal recorded during the 1-second time window spanning [-3.5, -2.5] seconds relative to the Go signal (or [-1.5, -0.5] with respect to the Cue signal). Baseline normalization was performed by subtracting the mean and dividing by the standard deviation of the baseline. To improve robustness, the baseline metrics were computed as the average across all trials within a session.

### Neural subspaces identification

The methods used to identify neural subspaces are based on a slight adaptation of those presented by Dekleva and colleagues [14].

The identification of neural subspaces was performed with trial-averaged data for each condition. Specifically, we constructed two data matrices, *Data_Ex_*and *Data_Im_*, both with dimensions *MT* ×*D_tot_*, where *M* is the number of concatenated movements, *T* is the number of time points (400, corresponding to 4 seconds of movement), and *D_tot_* is the total number of features (224, corresponding to 32 channels × 7 frequency bands). These matrices encode the full neural trajectories in the original recording space for execution and imagination conditions. Searching for neural subspaces within this space involves identifying covariance patterns across channels and frequency bands.

The first step was to identify a common latent space by reducing the dimensionality of the original recording space. To ensure that both *Data_Ex_*and *Data_Im_* were well-represented in this latent space, we performed Principal Component Analysis (PCA) separately on each dataset. We retained components *D_Ex_* and *D_Im_*, corresponding to 99% of the explained variance for execution and imagination, respectively. To find a shared orthonormal basis spanning the two spaces, we concatenated the matrices, forming *W* ∈ R*^D^_tot×_*_(_*^D^_Ex_*_+_*^DI^_m_*_)_, and applied Singular Value Decomposition (SVD). The left singular vectors *L* ∈ R*^D^_tot×_^D^_L_* defined a latent space equally representing both conditions, where *D_L_* = *D_Ex_* + *D_Im_*. Neural activity in the original space was then projected onto this latent space, yielding *Data_Ex L_* and *Data_Im L_*.

Next, PCA was applied separately to *Data_Ex L_* and *Data_Im L_* to facilitate the identification of neural subspaces exclusive to execution or imagination. To isolate a subspace exclusive to imagination, we focused on the last principal components (PCs) from the PCA of *Data_Ex L_*. By construction, these PCs explained minimal variance in execution data while potentially capturing variance relevant to imagination. We selected *D_Ex−null_*, the number of PCs explaining less than 1% of execution variance, and formed the execution-null space *U_Ex−null_* ∈ R*^D^_L×_^D^_Ex−null_* . By projecting *Data_Im L_* onto *U_Ex−null_* and performing another PCA to retain the leading *D_Excl−Im_* components explaining 99% of the variance, we obtained *W_Excl−Im_* ∈ R*^D^_Ex−null×_^D^_Excl−Im_* . The product of *U_Ex−null_* and *W_Excl−Im_* defined a subspace containing variance exclusively relevant to imagination, *S_Excl−Im_* ∈ R*^D^_L×_^D^_Excl−Im_* . Projecting latent activity from both conditions onto this subspace yielded imagination-specific responses. A similar procedure was performed to identify the execution-exclusive subspace.

However, *S_Excl−Im_* and *S_Excl−Ex_* were not orthogonal because they were derived from independent PCAs performed on the common latent space. To refine these subspaces and ensure orthogonality, we used the Python toolbox Pymanopt to optimize the manifolds via gradient descent [48]. This process minimized the sum of squared errors between the exclusive responses derived earlier and those obtained by projecting onto subspaces *Q_excl−Ex_* and *Q_excl−Im_*, which were constrained to be orthogonal as part of the same orthonormal matrix belonging to a Stiefel manifold *M* ∈ R*^D^_L×_*_(_*^D^_excl−Ex_*_+_*^D^_Excl−Im_*_)_.

Finally, the shared subspace *Q_Sh_*, capturing variance common to both execution and imagination neural activity, was determined as the null space of *Q* = [*Q_excl−Ex_, Q_excl−Im_*], where the brackets indicate concatenation. The dimensionality of the latent space was thus decomposed as *D_L_* = *^D^_excl−Ex_* ^+^ *^D^_excl−Im_* ^+^ *^D^_Sh_*.

### Alignment index and controls

We quantified the similarity between two neural subspaces using the alignment index introduced by [11]. The alignment index, *A*, is defined as:

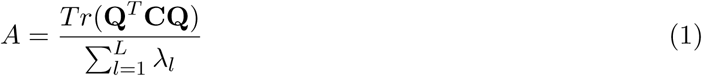

where *Q* represents the L-dimensional neural subspace onto which the data is projected, *C* the covariance matrix of the neural data, *λ_i_*is the *i^th^*largest eigenvalue of *C*, and *Tr*() is the trace of the matrix. The alignment index *A* quantifies the ratio of variance explained by the leading components within a subspace (principal components derived from the same activity) to the variance explained by an equivalent number of leading components derived from the opposing activity. This index ranges from 0 to 1, where 0 represents total orthogonality and 1 signifies complete alignment.

To assess the potential alignment observable by chance, we performed a random selection of neural subspaces following the procedure described in [10, 11]. Specifically, the random subspaces were sampled from a space defined by the real covariance matrix of the neural data. If the observed alignment index exceeds this chance-level control, it indicates that the two subspaces are more aligned than expected from random definitions.

Another critical control for alignment between subspaces involves estimating the level of alignment expected if the only source of misalignment was intrinsic variability across trials. To evaluate this, we computed the alignment index after shuffling the labels between Execution and Imagination data 1000 times, while maintaining the consistency of the movement label. An alignment index lower than this shuffling control suggests that the two spaces are less aligned than would be expected if there was no distinction between the conditions.

### Features contributions to neural subspaces

To understand the contributions of individual channels and frequency bands to a given neural space, we computed the overall transformation matrix that projects the original data into a specific subspace. This was achieved by multiplying the transformation matrix that projects data onto the common latent space with the matrix that projects from the latent space into the specific neural subspace. For example, for Execution data:

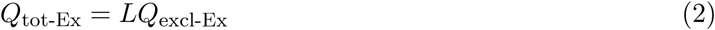

where *L* ∈ R*^D^*_tot_*_×DL_* is the transformation matrix projecting data into the latent space, and *Q*_excl-Ex_ ∈ R*^D^_L×_^D^*_excl-Ex_ is the matrix projecting from the latent space into the Execution-exclusive subspace. To quantify the contribution of each feature (i.e., combinations of channels and frequencies), we calculated the *L*_2_-norm of the *D*_excl-Ex_-dimensional vectors in *Q*_tot-Ex_. To account for variations in channel activity levels, we scaled these contributions by the amplitude of the average activity across trials, computed using concatenated Execution and Imagination data. This yielded a vector **C**_Ex_ ∈ R*^D^*_tot_*_×_*_1_, where each entry represents the contribution of a specific channel-frequency combination.

Finally, contributions were normalized within each neural space to express them as percentages:

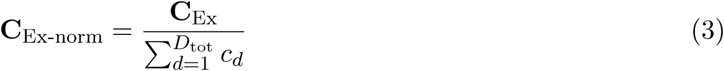

where *c_d_* represents the contribution of the *d*-th feature. This normalization provided the percentage contribution of each feature to the specific neural space.

### Movement decoding

The data, originally structured as *N*_trials_ × *T* × *D*_tot_ (where *T* represents the number of timestamps and *D*_tot_ is the number of dimensions in the recording space), were first projected onto the neural space of interest. Subsequently, the data were segmented into non-overlapping time windows of 50 milliseconds, with the average computed within each window. This process yielded *N*_tw_ time windows, where *N*_tw_ is the total number of extracted windows. To create feature vectors, the temporal evolution for each dimension was flattened by concatenating the data across all time windows, resulting in a final structure of *N*_trials_ × [*N*_tw_ × *D*_space_], where *D*_space_ denotes the dimensionality of the neural space of interest.

To reduce the dimensionality of the dataset, PCA was applied to the feature matrix, retaining 99% of the variance. PCA was fitted on the training set and subsequently applied to the test set. A Linear Discriminant Analysis (LDA) classifier was then employed, and performance was evaluated using balanced accuracy, defined as the average recall across all classes. When specified, a 10-fold stratified cross-validation was used. For generalization analyses, such as the ones in **Fig. 5** and **Fig. 6F**, a bootstrapping method was used, where trials from the training and test sets where randomly resampled 1000 times. To assess the statistical significance of the results, permutation tests were performed by shuffling the movement labels 1000 times and recalculating the metrics for each shuffle.

## Acknowledgments

The authors sincerely thank the participant for his invaluable time, dedication, and commitment throughout the data collection process. We also extend our gratitude to the multidisciplinary technical and clinical teams at Clinatec (CEA-LETI and CHU-Grenoble Alpes) for their involvement in the BCI&Tetraplegia clinical trial (NCT02550522), which enabled the advancements presented in this work. Finally, we thank Vincent Auboiroux for his assistance in providing medical reconstruction images of the implant location.

S.M. and E.R. were supported by the #NEXTGENERATIONEU (NGEU) and funded by the Ministry of University and Research (MUR), National Recovery and Resilience Plan (NRRP), project: MNESYS (PE0000006) – A Multiscale integrated approach to the study of the nervous system in health and disease (DN. 1553 11.10.2022). S.M. also received support from two additional NRRP projects: THE (IECS00000017) – Tuscany Health Ecosystem (DN. 1553 11.10.2022), and BRIEF (IR0000036) – Biorobotics Research and Innovation Engineering Facilities (DN. 103 17.06.2022). L.P., V.d.S., S.M., and S.S. were supported by the Bertarelli foundation and the Swiss National Science Foundation (Grant Number 10003473). L.S., S.K., S.C., T.A., and G.C. were supported by the CEA, the Carnot Institute CEA-Leti, the French National Research Agency (ANR), the French Ministry of Health and Research and the Fonds Clinatec.

## Authors contribution

L.P., S.S., and S.M. conceptualized the study. L.P., S.S., S.M., L.S., S.K., T.A., and G.C. designed the task and implemented the experimental setup. L.P., L.S., and V.d.S. collected the data. L.P., L.S., V.d.S., and E.R. analyzed the data. All authors participated in the interpretation of the results. L.P., S.S., and S.M. wrote the manuscript with input from all authors.

