## Supplementary Material for "Decoupling simultaneous motor imagination and execution via orthogonal ECoG neural representations"

#### Supplementary Figures

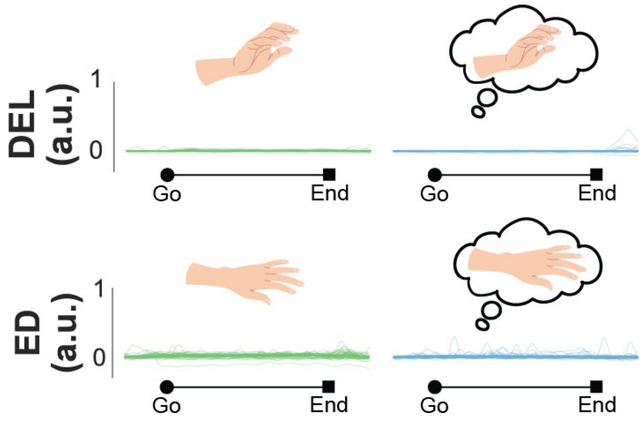


**Figure S1: EMG activity of non-target muscles of each task for one example session.** Top: Lateral deltoid activity during the wrist extension movement. Bottom: Extensor digitorum activity during the reaching movement. Execution trials are represented in green, while imagination trials are represented in blue.

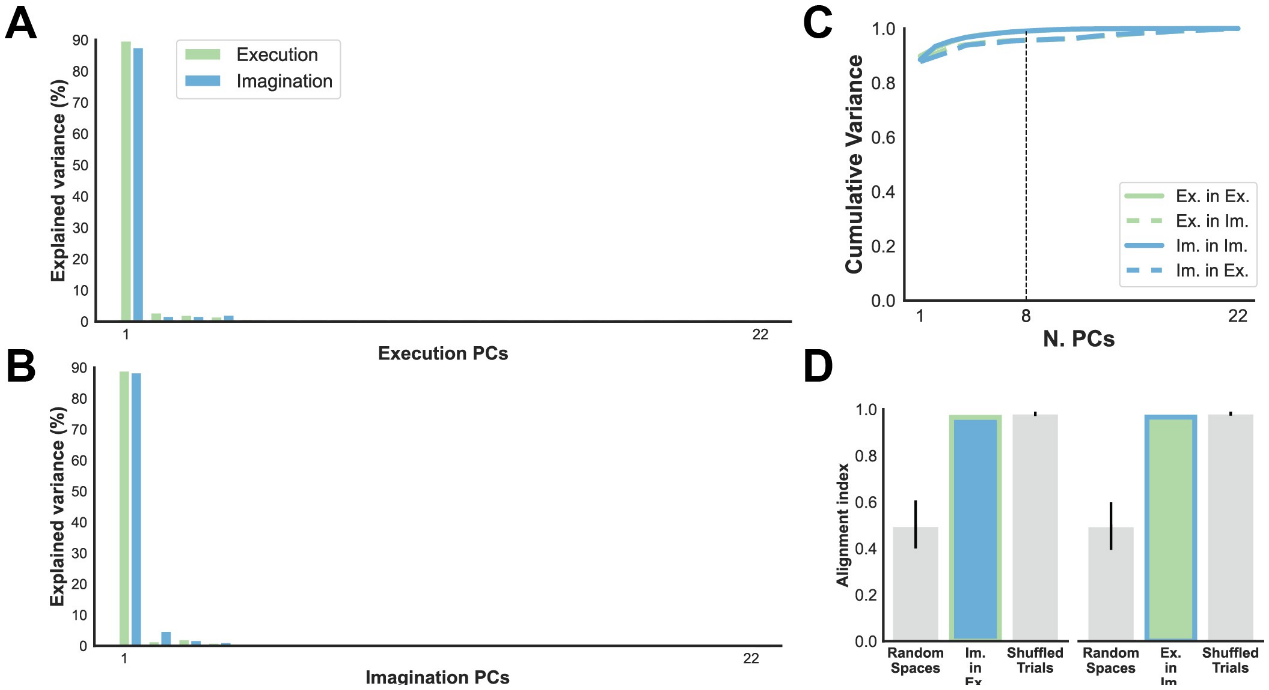


**Figure S2: Neural data from the intertrial period is completely aligned. A)** Variance explained by all PCs when projecting execution and imagination intertrial data onto the PCs computed over the execution data. **B)** Same as **A** but for PCs computed over imagination data. **C)** Cumulative variance for the projection within and across spaces. **D)** Alignment indices for both cross-projections together with random and labels-shuffling controls.

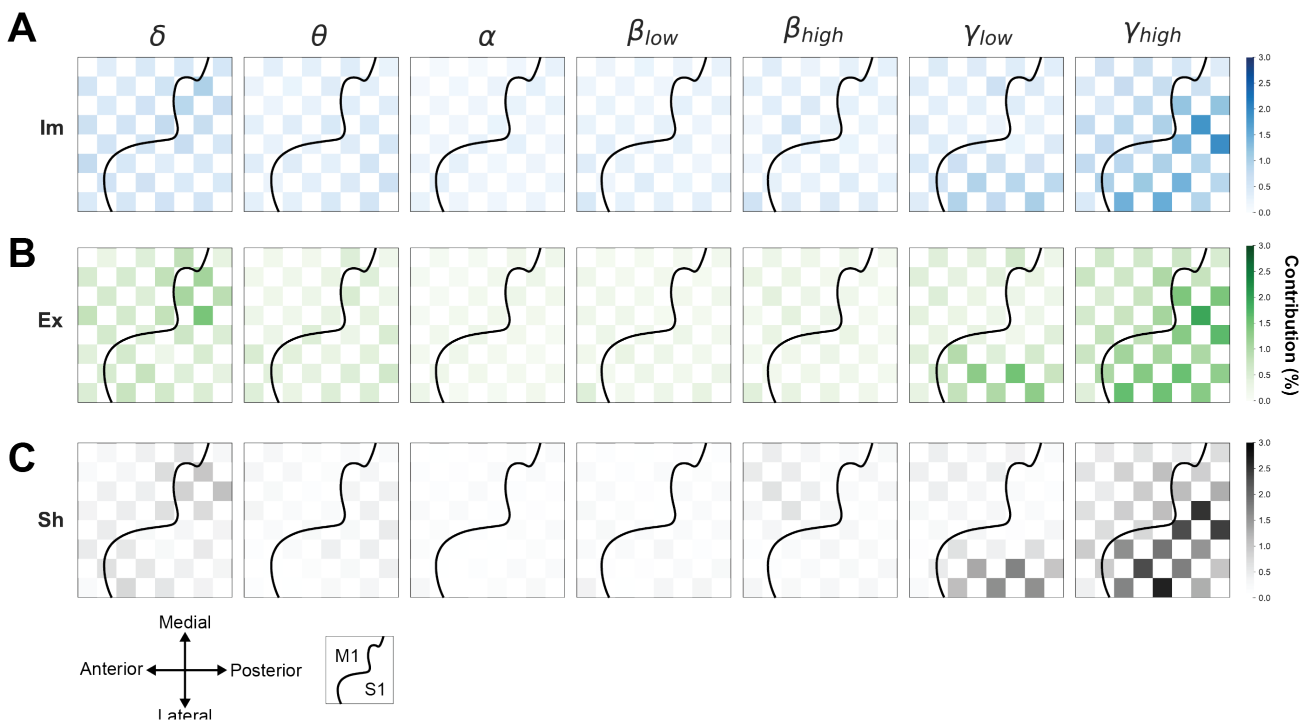


**Figure S3: Distribution of channel contributions over the electrode grid.** Contributions normalized per space and divided per band for **A)** Imagination space, **B)** Execution space, and **C)** Shared space. The curved black line represents the approximate location of the central sulcus.

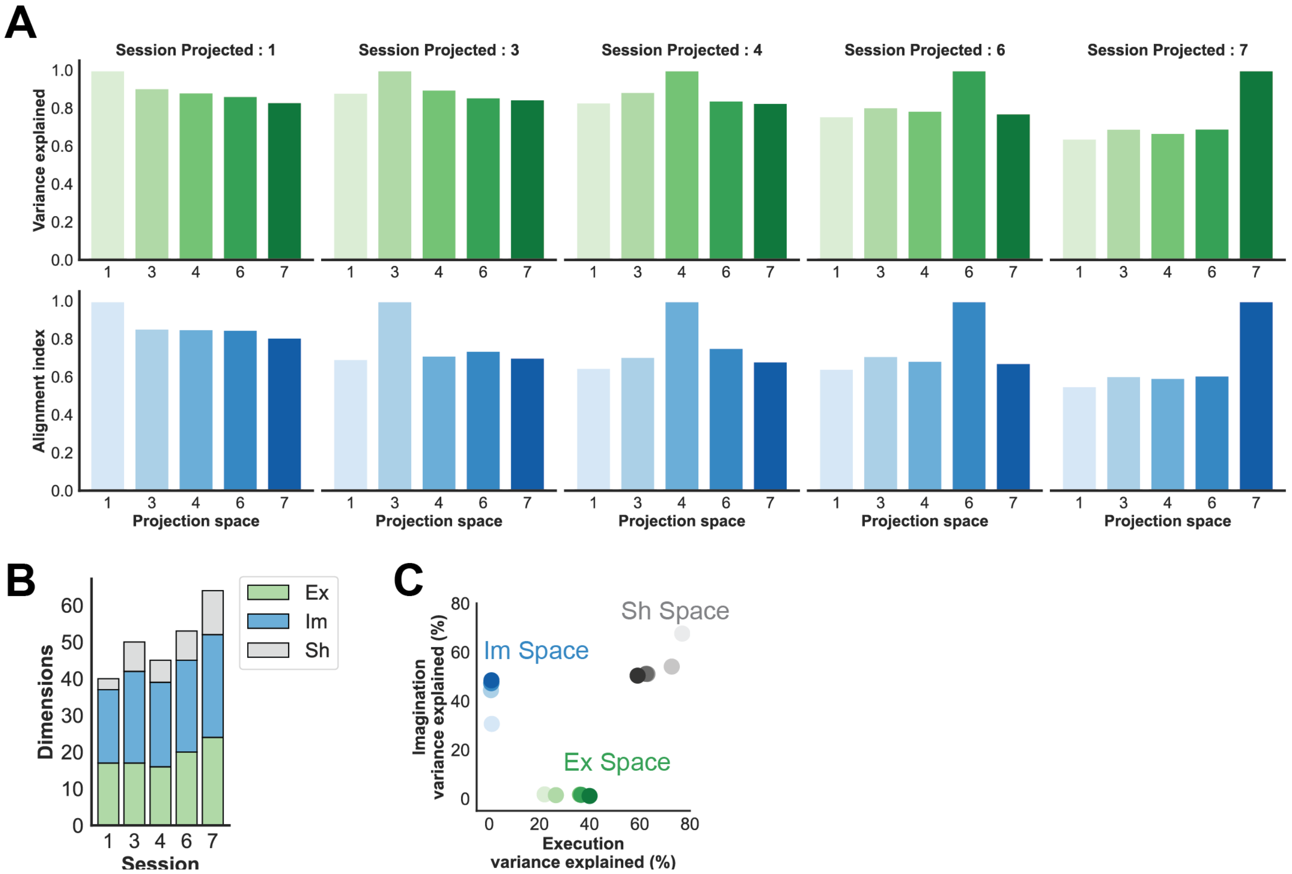


**Figure S4: Variance stability across individual sessions. A)** Variance explained for every session (columns) for both execution (top row) and imagination (bottom row) data when projecting data onto the latent spaces identified on individual sessions. **B)** Dimensionalities of the neural spaces identified on the data of individual sessions. **C)** Variance explained by the individual neural spaces for each session separately.

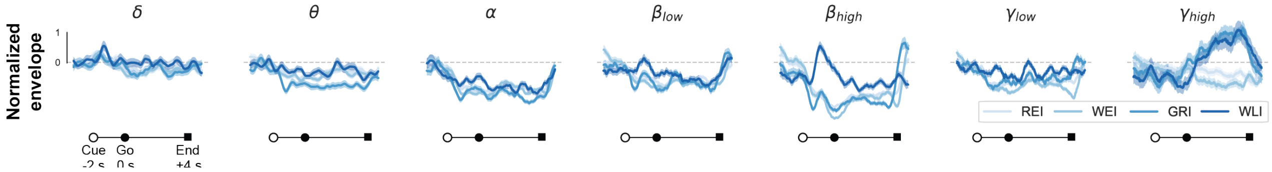


**Figure S5: Example neural signals for the 4 imagined movements**. Neural envelopes for all frequency bands are shown for the same example channel located on M1 presented in Fig.1H. Thick lines represent the average across trials, while the shaded area indicates the standard error of the mean.

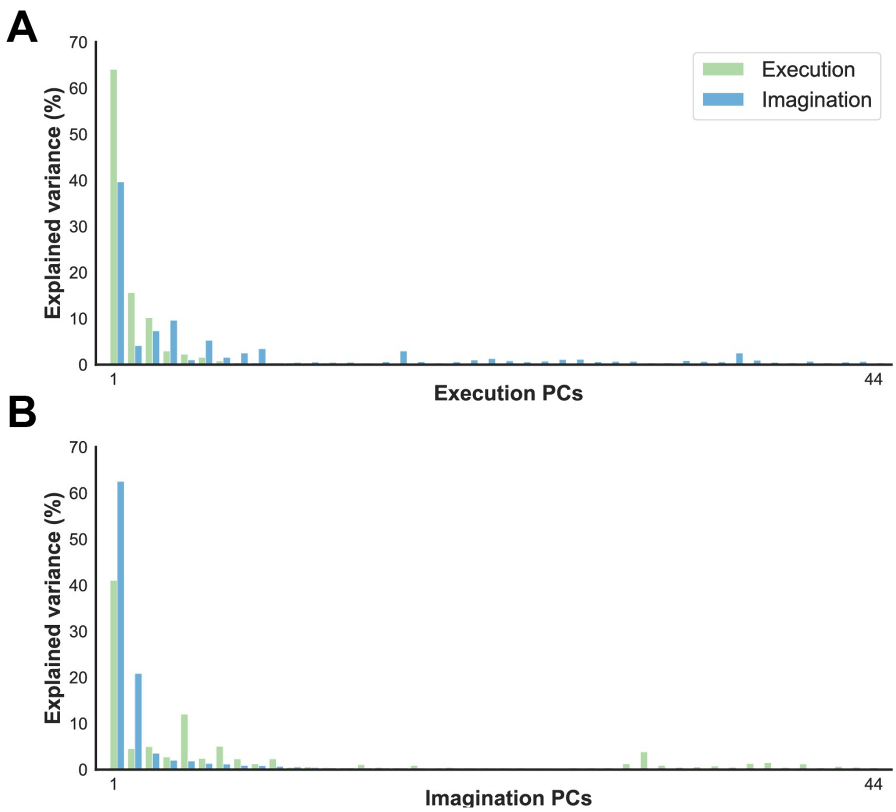


**Figure S6: Variance explained by each PC when considering four imagined movements**. PCA performed on **A)** execution activity, **B)** imagination activity.

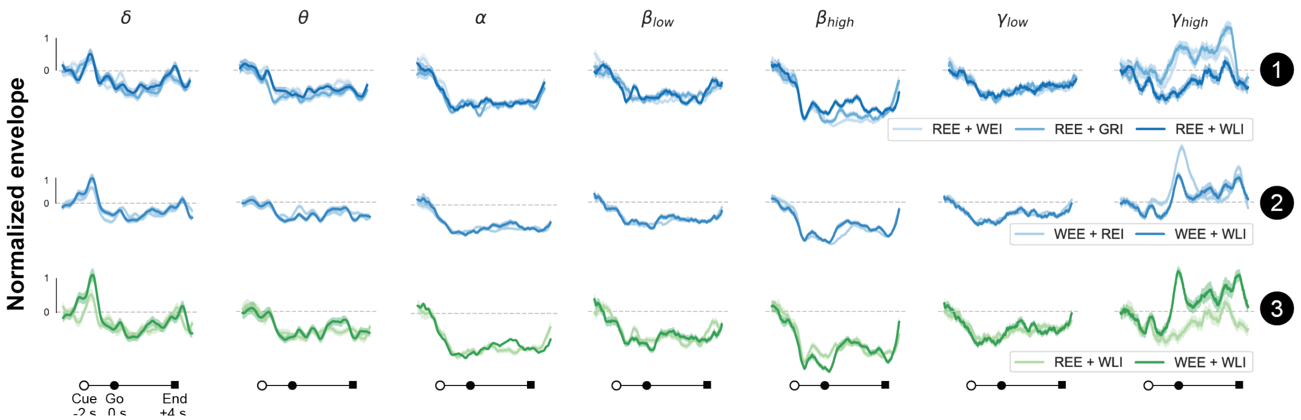


**Figure S7: Example neural signals for the dual task scenario**. Neural envelopes for all frequency bands are shown for the same example channel located on M1 presented in Fig.1H. 1, 2, and 3 refer to the possible combinations of movements during the dual task paradigm: 1 = executed reaching, 2 = executed wrist extension, 3 = imagined walking. Thick lines represent the average across trials, while the shaded area indicates the standard error of the mean.

### Supplementary Tables

**Table S1:** Dates of acquisition of the sessions recorded for this study.

| Session | Date | Days passed from session 1 |
|---|---|---|
| **1** | 27 Apr 2023 | 0 |
| **2** | 28 Apr 2023 | 1 |
| **3** | 05 Jun 2023 | 39 |
| **4** | 30 Jan 2024 | 278 |
| **5** | 31 Jan 2024 | 279 |
| **6** | 07 Mar 2024 | 315 |
| **7** | 08 Jul 2024 | 438 |
| **8** | 30 Sep 2024 | 522 |

**Table S2:** Overview of the experimental paradigm. Columns with the same color represent trials within the same blocks. Numbers in parentheses () indicate the number of blocks designated as single-task or dual-task.

| Session | Reaching Executed (REE) | Reaching Imagined (REI) | Wrist Extension Executed (WEE) | Wrist Extension Imagined (WEI) | Grasping Imagined (GRI) | Walking Imagined (WEI) | Single-tasks | Dual-tasks | EMG recorded | N. blocks |
|---|---|---|---|---|---|---|---|---|---|---|
| **1** | X | X | X | X | | | X | | | 16 |
| **2** | X | X | X | X | | | X (2) | X (8) | | 10 |
| **3** | X | X | X | X | | | X (6) | X (3) | | 9 |
| **4** | X | X | X | X | X | X | X | | X | 12 |
| **5** | X | X | X | X | X | X | | X | X | 12 |
| **6** | X | X | X | X | X | X | X | | X | 12 |
| **7** | X | X | X | X | X | X | X | | X | 9 |
| **8** | X | X | X | X | X | X | | X | X | 12 |

**Table S3:** Number of trials conducted per session and condition. The number in parentheses () represents the identified artifacts, while the number in brackets [] shows the percentage of trials retained relative to the total.

| Session | REE | WEE | REI | WEI | GRI | WLI | REE + WEI | REE + WLI | REE + WEE | REE + GRI | WEE + REI | WEE + WLI |
|---|---|---|---|---|---|---|---|---|---|---|---|---|
| **1** | 80 | 80 (1) | 80 | 80 | / | / | / | / | / | / | / | / |
| **2** | 10 | 10 (1) | 10 (1) | 10 (1) | / | / | 80 (1) | / | 80 (2) | / | 80 | / |
| **3** | 30 (1) | 30 (1) | 30 | 30 | / | / | 30 (1) | / | 30 (3) | / | 30 (2) | / |
| **4** | 60 | 60 (15) | 60 | 60 (10) | 60 (13) | 60 (17) | / | / | / | / | / | / |
| **5** | / | / | / | / | / | / | 72 (9) | 72 (5) | / | 72 (9) | 72 (1) | 72 (1) |
| **6** | 60 (8) | 60 (1) | 60 (4) | 60 (1) | 60 (3) | 60 (1) | / | / | / | / | / | / |
| **7** | 45 (10) | 45 | 45 (2) | 45 | 45 | 45 | / | / | / | / | / | / |
| **8** | / | / | / | / | / | / | 72 (3) | 72 (1) | / | 72 (1) | 72 | 72 (3) |
| **Total** | 266 [93%] | 266 [93%] | 278 [98%] | 273 [96%] | 149 [90%] | 147 [89%] | 240 [94%] | 138 [96%] | 105 [95%] | 134 [93%] | 251 [99%] | 140 [97%] |

**Table S4:** Additional results for movement decoding during dual-tasks as in Fig.6F with varying parameters. The grayed-out line highlights the results presented in the main text.

| | | **Reaching executed** | | **Wrist extension executed** | | **Walking imagined** | |
|---|---|---|---|---|---|---|---|
| **Time window duration** | **PCA variance retained** | **Excl. Im. space** | **Shared space** | **Excl. Im. space** | **Shared space** | **Excl. Ex. space** | **Shared space** |
| 50 ms | 0.99 | 42.8 ± 2.6 | 44.1 ± 2.5 | 51.8 ± 2.1 | 53.4 ± 1.2 | 68.3 ± 3.0 | 78.1 ± 2.6 |
| 50 ms | 0.95 | 44.5 ± 2.5 | 44.9 ± 2.6 | 52.2 ± 2.1 | 52.8 ± 1.2 | 69.8 ± 2.9 | 77.0 ± 2.5 |
| 50 ms | 0.9 | 45.4 ± 2.7 | 44.1 ± 2.6 | 52.1 ± 1.8 | 52.3 ± 1.0 | 72.0 ± 3.0 | 74.7 ± 2.6 |
| 50 ms | 0.8 | 43.8 ± 2.4 | 44.9 ± 2.9 | 53.8 ± 1.9 | 54.1 ± 1.3 | 74.3 ± 3.0 | 72.4 ± 3.0 |
| 250 ms | 0.99 | 43.3 ± 2.7 | 44.6 ± 2.7 | 51.6 ± 2.4 | 52.8 ± 1.2 | 69.4 ± 3.1 | 77.7 ± 2.7 |
| 250 ms | 0.95 | 44.8 ± 2.6 | 45.0 ± 2.4 | 52.0 ± 2.0 | 52.3 ± 1.0 | 71.9 ± 3.1 | 76.0 ± 2.5 |
| 250 ms | 0.9 | 45.0 ± 2.6 | 44.1 ± 2.5 | 52.1 ± 1.9 | 52.5 ± 1.1 | 72.9 ± 2.9 | 74.0 ± 2.5 |
| 250 ms | 0.8 | 44.0 ± 2.5 | 49.0 ± 3.3 | 53.9 ± 2.0 | 54.5 ± 1.5 | 74.9 ± 2.9 | 69.6 ± 2.5 |
